# Comprehensive proteomic and metabolomic profiling of *mcr-1* mediated colistin resistance in *Escherichia coli*

**DOI:** 10.1101/434548

**Authors:** Hui Li, Yingyu Wang, Qingshi Meng, Yang Wang, Guoliang Xia, Xi Xia, Jianzhong Shen

## Abstract

The spread of *mcr-1* in human and veterinary medicine has jeopardized the use of polymyxins, the last-resort antibiotics against life-threatening multidrug-resistant Gram-negative bacteria. As a lipid-modified gene, whether *mcr-1* brings proteomic and metabolomic changes in the bacteria and affects the corresponding metabolic pathway is largely unknown. Herein, we used label-free quantitative proteomics and untargeted metabolomics to profile comprehensive proteome and metabolome characteristics of *mcr-1*-mediated colistin-resistant and -sensitive *Escherichia coli* and further insight the resistant mechanism of colistin. We identified large sets of differential expression proteins and metabolites that contributed to *mcr-1*-mediated antibiotic resistance predominantly in the different growth conditions with and without colistin. *mcr-1* could cause the down-regulated expression of most proteins to adapt drug pressure. Pathway analysis showed that metabolic process was significantly affected, mainly related to glycerophospholipid metabolism, thiamine metabolism, and lipopolysaccharide biosynthesis. The substrate phosphatidylethanolamine for *mcr-1* to mediate colistin resistance is accumulated in colistin-resistant *E. coli*. Notably, *mcr-1* can not only cause the phosphoethanolamine modification of bacterial cell membrane lipid A, but also affect the biosynthesis and transport of lipoprotein in colistin resistance through disturbing the expression of efflux pump proteins involved in cationic antibacterial peptide resistance pathway. Overall, the disturbed glycerophospholipid metabolism, lipopolysaccharide biosynthesis and the accumulation of the substrate phosphatidylethanolamine is closely related with *mcr-1*-mediated colistin resistance and these findings can further provide valuable information to inhibit colistin resistance by blocking this metabolic process.

## Author summary

Since the discovery of the plasmid-mediated colistin resistance determinant *mcr-1* in Enterobacteriace in 2015, its widespread dissemination brings potential threats to the treatment of extensively drug resistant bacterial infections in animals and humans. Although many excellent achievements in its epidemic evidence and resistance mechanism have been made, the in-depth colistin resistance mechanisms, such as the omics profile of bacteria mediated by *mcr-1* in *Escherichia coli* needs further research. In this study, comprehensive proteomic and metabolomic profiles of *mcr-1*-mediated colistin-resistant and -sensitive *Escherichia coli* were investigated to clarify the protein and metabolite features related to *mcr-1*. The proteome characteristics of colistin-resistant *E. coli* carrying *mcr-1* under different selection pressures of colistin and polymyxin B were investigated, and the metabolic pathways including glycerophospholipid metabolism, thiamine metabolism, and lipopolysaccharide biosynthesis were disturbed. Up to 67 differential metabolites were identified in *mcr-1*-mediated colistin-resistant strains, demonstrating changes in polyketide sugar unit biosynthesis, glycerophospholipid metabolism, and phosphatidylethanolamine biosynthesis. Disturbed glycerophospholipid metabolism, lipopolysaccharide biosynthesis and the accumulation of the substrate phosphatidylethanolamine for *mcr-1* were the typical changes in *mcr-1*-mediated colistin resistance in *E. coli*. We expect that this proteome and metabolome profiling data set illustrates the in-depth *mcr-1*-mediated colistin resistance mechanisms from an omics perspective.

## Introduction

Resistance to polymyxins has emerged worldwide threatening the efficacy of the last-resort antimicrobials used for the treatment of multidrug-resistant (MDR) *Enterobacteriaceae* infection in humans [1, 2]. In parallel, the heavy use of colistin in veterinary medicine is currently being reconsidered due to increased reports of colistin-resistant bacteria and has been banned as feed additives in China in 2016 [3]. Colistin resistance can be mediated by phosphoethanolamine (PEA) or 4-amino-arabinose modification of lipid A that abolishes the initial electrostatic interaction with polymyxins [4, 5]. Colistin resistance was appeared in MDR bacterial isolates with chromosomal resistance mutations in lipid A biosynthesis and regulation and, more recently, mobilized colistin resistance determinants *mcr-1 to −8* [6–13]. *mcr-1* was the firstly reported plasmid-mediated colistin resistance gene and encoded a member of the family of PEA transferases that decorated the lipid A headgroups of lipopolysaccharide (LPS) with PEA [6]. The resistance rate of *mcr-1* increased rapidly, and the gene has the risk of entering the human body from the food chain, which is a serious threat to human health and public health security [14, 15].

Since the *mcr-1* gene was reported, many studies clearly clarified the status quo of its transmission and provided its epidemic evidence in food, livestock and human [16–18]. How to effectively avoid the public health hazards induced by *mcr-1* has become an important challenge. In an effort to encompass these matters and increase the metabolic coverage of available methodologies, system-level approaches have applied in unknown functional discovery. Proteomics and metabolomics, as a new “omic” technique, can study the whole protein expression and the dynamic changes of metabolites in specific tissues or cells on a system level, and elucidate the developmental process of specific biological processes and regulatory mechanism at the molecular level [19, 20]. In addition, the use of proteomics techniques to study bacterial resistance mechanisms made it possible to find potential large–scale targets for antimicrobials [21, 22]. Fernandez-Reyes *et al* [23] studied the resistant cost of colistin-resistant *Acinetobacter baumannii* and found that outer membrane proteins, chaperones, protein biosynthesis factors and metabolic enzymes were down-regulated in colistin-resistant strains from a proteomic perspective. It was noteworthy that the fatty acyl side chains with the C_12_-C_14_ carbon chain of the lipid A group, which was essential for the maintenance of bacterial LPS function, were involved in the synthesis of bacterial fatty acid type **II** synthesis pathway, suggesting that it was essential to decipher the metabolic mechanism of *mcr-1* mediated resistance from a metabolic perspective. Hua *et al* [24] utilized a multiomics approach to explore the response of multiple drug-resistant *A. baumannii* to colistin and found that RND efflux pump system were over-expressed and inhibited the expression of FabZ and β-lactamase. However, as a lipid-modified gene, the comprehensive proteomic and metabolomic profiling of *mcr-1*-mediated colistin resistance in bacteria is still unknown.

In this study, we investigated the comprehensive proteome and metabolome changes in *mcr-1*-mediated colistin-resistant and -sensitive *E. coli* strains using label-free quantitative proteomics and untargeted metabolomics. The differential expressed proteins (DEPs) and metabolites were successfully selected to decipher the in-depth colistin resistance mechanisms mediated by *mcr-1*. This study could provide insight into the molecular mechanisms and potential disturbed metabolic pathways of *mcr-1* mediated bacterial resistance.

## Materials and Methods

### Strains, plasmid construction and antibiotic susceptibility testing

The gene encoding *mcr-1* with its own promoter (156 bp) and downstream sequence (111 bp) was amplified by PCR from the plasmid pHNSHP45 [6] using Q5 High-Fidelity DNA Polymerase (New England Biolabs, Ipswich, USA) and cloned into the plasmid pUC19 using Eco **I** and Xba **I** sites. Then the constructed plasmid pUC19-*mcr-1* was transformed into *E. coli* component cells DH5α. Transformants were selected by the inclusion of ampicillin (100 *μ*g/mL) on Luria Broth (LB) agar plates. Cloning results were confirmed by PCR sequencing. The successfully constructed strains *E. coli* DH5α (pUC19-*mcr-1*) carrying an 1,893 bp fragment were used as the engineering strains. The blank plasmid pUC19 was chemically transformed into *E. coli* DH5α and considered as the control strains (*E. coli* DH5α (pUC19)). Susceptibility testing was carried out on the above constructed *mcr-1*-expressing strains and control strains against polymyxin B and colistin according to the Clinical and Laboratory Standards Institute guidelines [25]. The minimum inhibitory concentrations (MICs) of *E. coli* DH5α (pUC19-*mcr-1*) for colistin and polymyxin B was 4 *μ*g/mL, an 8-fold increase compared with the control strains.

### Protein lysis and digestion

Sample of colistin-resistant *E. coli* DH5α (pUC19-*mcr-1*) strains for label-free quantitative proteome analysis was taken from cells that were grown on LB broth supplemented with colistin and polymyxin B under three different drug culture conditions (0, 0.5, and 1 *μ*g/mL), respectively. And colistin-sensitive *E. coli* DH5α (pUC19) strains were cultured in blank LB broth at 37 °C overnight until they reached in exponential state. Three biological replicates were performed for each condition. Cells were centrifugated at 10,000 × g, 4 °C for 30 min and washed twice with 10 mL ice-cold PBS buffer, and stored at −80 °C until further processing. Cells were re-suspended in 1 mL lysis buffer (8 M urea, 50 mM Tris-HCl (pH 7.4), 0.5%SDS, 1 mM PMSF, 20 *μ*L protease inhibitor cocktail). The cells were lysised by indirect sonication (35% energy, 4 min) in a Sonics VCX105 (Sonics & Materials, Inc., Newtown, USA) and centrifugated at 20,000× g, 4 °C for 30 min. A small aliquot of the supernatant was taken to determine the protein concentration using a Pierce BCA Protein Assay Kit (Thermo Fisher Scientific, Rockford, USA). All protein samples were normalized to the same concentration with cell lysate, using the protein sample with the lowest concentration as a reference. A total of 100 *μ*g protein were taken and added 0.1 M triethylammonium bicarbonate (TEAB) to a final volume of 100 *μ*L. Protein were reduced with a final concentration of 50 mM dithiothreitol buffer at 37 ° for 1 h and alkylated with iodoacetamide (100 mM final concentration) in the dark for 30 min. Five volumes of cold acetone (-20 °C) were added for protein precipitation and the solution was incubated overnight at −20 °C. After centrifugation at 20,000 × g for 20 min, the precipitate was stayed at room temperature for 2-3 min until acetone volatilization. 100 *μ*L of 0.1 M TEAB was added to dissolve the protein sample. Trypsin (2 *μ*g) in 0.1 M TEAB were added to digest protein and incubated at 37 °C for 16 h. After digestion, the sample was evacuated in Speedvac rotation evaporator (Thermo Fisher Scientific, Japan). The solution was reconstituted with 100 *μ*L of water/acetonitrile (98/2, v/v, containing 0.1% formic acid) and drained again until almost no white salts were visible in the microcentrifuge tube. Peptides were dissolved in 100 *μ*L of water/acetonitrile (98/2, v/v, containing 0.1% formic acid) and centrifuged at 20,000 × g for 15 min. The supernatant was collected and injected onto the NanoLC-MS analysis.

### Label-free quantitative proteome analysis

The label-free quantitative analysis was performed by Eksigent 2D coupled to a TripleTOF 6600 mass spectrometer (Sciex, Framingham, MA, USA). Peptides were first loaded on a trap column 120Å ChromXP C_18_-CL resin (200 *μ*m×0.5 mm, 3 *μ*m), and then separated on a nano column Eksigent cHiPLC column 120Å ChromXP C_18_-CL resin (75 *μ*m×15 cm, 3 *μ*m). The flow rate was 300 nL/min over a 95 min multi-segment gradient on mobile phase A (water:dimethyl sulphoxide:acetonitrile, 96:2:2, v/v/v, containing 0.1% formic acid) and mobile phase B (acetonitrile:dimethyl sulphoxide:water, 96:2:2, v/v/v, containing 0.1% formic acid). The gradient conditions were optimized as follows: 0-50 min, 5%-20% B; 50-70 min, 20%-32% B; 70-75 min, 32%-80% B; 75-80 min, 80% B; 80-95 min, 80%-5% B.

For data dependent acquisition (DDA), a TOF-MS scan over a mass range *m/z* 350-1500 Da with 250 ms accumulation time, followed by *m/z* 100-1800 Da for MS/MS scans in high sensitivity mode with 80 ms accumulation time of up to TOF 40 ion candidates per cycle was performed. For data independent acquisition (DIA), a set of overlapping 25 transmission windows, each 25 Da wide, was constructed and covered the precursor mass range of *m/z* 400-1000 Da. The SWATH product ion scans were acquired in the range of *m/z* 100-1800 Da. The SWATH-MS1 survey scan was acquired with the accumulation time of 50 ms, and it was followed by 40 ms accumulation time high sensitivity product ion scans.

Data analysis was performed using Skyline [26]. ProteinPilot v5.0 was utilized to carry out spectral counting for label-free quantification. DEPs were filtered by the following cutoff: false discovery rate < 1% and peptide confidence threshold > 99%, fold change > 2.0 and *p*-value < 0.01.

### Quantitative RT-PCR

A total of fourteen DEPs (*tolA*, *yqaB*, *nadC*, *ybiG*, *htp*, *glpQ*, *osmY*, *waaJ*, *yaaF*, *eutE*, *ydcH*, *yjiY*, *gutD*, and *yccX*) were verified by qRT-PCR using the iQ SYBR Green Super Mix (Qiagen) and analysed by a 7500 Real-Time PCR system (Applied Biosystems, Thermo Fisher Scientific, Inc.). Primer sequences were included in Table S1. Relative expression of these selected genes was normalized to the expression of 16S rRNA gene. Amplification efficiency and relative transcript abundance were analyzed using 2^-ΔΔCT^ method and *t*-tests.

### LC/MS metabolomic analysis

The colistin-sensitive *E. coli* DH5α (pUC19) strains were cultured in blank LB broth at 37 °C overnight, and colistin-resistant *E. coli* DH5α (pUC19-*mcr-1*) strains were grown on LB medium under three different culture concentrations of colistin (0, 0.5, and 1 *μ*g/mL). The bacterial turbidity was adjusted to OD_600_ of 0.8 using blank sterile LB broth. 10 mL aliquots were directly transferred to 50 mL tubes and pelleted by fast centrifugation (5 min at 10,000 × g, 0 °C). The supernatant was discarded, and the pellets were flash frozen in liquid nitrogen to quench the cell metabolism. Subsequently, cell pellet was extracted with 1.5 mL methanol at −20 °C and vortexed for 1 min. The freeze-thawing operations in liquid nitrogen were repeated three times to lysis the cells. The samples were centrifuged at 12,000 × g for 15 min at −10 °C (Beckman Coulter Inc., USA) and re-extracted the cellular metabolites again using the above procedure. The cellular extracts were combined and dried under vacuum using a SpeedVac V100 (Thermo Fisher Scientific, Inc.). The extracted metabolites were re-suspended in 500 *μ*L acetronile:water (50:50, v/v, containing 0.1% formic acid). A quality control (QC) sample was prepared by mixing aliquots from all supernatant samples (50 *μ*L from each sample). Mixed standard solution (sulfadiazine, difloxacin, monensin, tylosin, and thiamphenicol) at a final concentration of 200 ng/mL was spiked into the extracts for quality control (QC) purposes of compound mass.

The cellular metabolites were analyzed by a Waters ACQUITY ultra-high performance liquid chromatography (UPLC) coupled with a hybrid TOF-MS SYNAPT HDMS (Waters, Manchester, U.K.). The weakly polar metabolites were separated by BEH Shield RP C_18_ column (50 × 2.1 mm, 1.7 *μ*m) with mobile phase A (0.1% formic acid in water) and mobile phase B (0.1% formic acid in acetonitrile) at a flow rate of 0.3 mL/min at 35 °C. The optimized gradient elution procedure was as follows: 0-1 min, 2-5% B; 1-8 min, 5-90% B; 8-10 min, 90-100% B; 10-12 min, 100-2% B. The strongly polar metabolites were analyzed by BEH HILIC column (50 × 2.1 mm, 1.7 *μ*m) and eluted using mobile phase A (acetonitrile:water, 95:5, v/v, containing 0.05 mM ammonium acetate) and mobile phase B (acetonitrile:water, 50:50, v/v, containing 0.05 mM ammonium acetate) at a flow rate of 0.3 mL/min at 35 °C. The gradient elution conditions were optimized as follows: 0-1 min, 1% B; 1-2 min, 1-10% B; 2-4 min, 10% B; 4-5 min, 10-30% B; 7-8.5 min, 30-50% B; 8.5-10min, 50-100% B; 10-12 min, 100-1% B. QTOF/MS spectrum was operated in ESI+ and ESI^−^ modes, respectively. The mass range was set at *m/z* 50-1200 Da in the full-scan mode. The optimized ESI parameters were as follows: capillary voltage, 3.0 kV; cone voltage, 35 V, source temperature, 100 °C, desolvation temperature, 450 °C, desolvation gas flow, 600 L/h. For accurate mass measurement, leucine enkephalin was used as the lock spray standard ([M+H]+ =556.2771; [M-H]- =554.2615) at a concentration of 100 ng/mL with a flow rate of 50 *μ*L/min.

The datasets were processed in Progenesis QI software v. 2.3 (Nonlinear Dynamics, Newcastle, UK) using the following parameters: the automatic sensitivity method value was set at 1 (fewer), peak widths at and less than 0.19 min were ignored, ions after 11 min were ignored, and each sample was normalized to the QC sample. The multivariate analysis, including principal component analysis (PCA) and partial least squares-discriminant analysis (PLS-DA), was constructed to determine the distributions and find the metabolic differences between the control strains and the resistant strains. The filtered data (*p* < 0.05 and fold change ≥ 2) were processed for further analysis and display. The assigned modified metabolite ions were identified by database searches in the ECMDB (http://www.hmdb.ca/spectra/ms/search), Chemspider (http://www.chemspider.com/), HMDB (http://www.hmdb.ca/), and Metlin (https://metlin.scripps.edu/) database. The mass tolerance for the database search was set at 10 ppm.

### Data analysis

Metabolic pathway of DEPs and differential metabolites were analyzed using KEGG software (http://www.genome.jp/kegg/) to investigate the disturbed metabolic pathways and facilitate biological interpretation. The DEPs, metabolites and corresponding pathways were imported into Cytoscape software (v.3.4.0) for visualization of the network models. The heat map was constructed with R software (v. 3.4.4) to show the key information related with comprehensive changes induced by *mcr-1* gene in colistin resistance.

## Results

### Comprehensive proteome changes under different drug culture conditions

To define proteome changes accompanying *mcr-1* mediated colistin resistance, we compared protein levels of colistin-resistant and -sensitive *E. coli* strains in different culture conditions by label-free quantitative mass spectrometry using SWATH-MS acquisition mode. A total of 2,784 proteins were identified, which represented approximately 64.3% coverage of the predicted proteome. The protein distribution detected in colistin-resistant *E. coli* DH5α (pUC19-*mcr-1*) proteome using SWATH-MS was shown in Fig 1A. A total of 57 proteins (26 up-regulated and 31 down-regulated) were differentially expressed when no drug supplemented in the media (Fig 1B and Table S2). When the colistin-resistant *E. coli* DH5α (pUC19-*mcr-1*) strains were grown under different selection pressure of colistin and polymyxin B (0.5 and 1 *μ*g/mL), the number of DEPs was significantly increased and most of the proteins were down-regulated (Fig 1B and Table S3-S6). The resistant *E. coli* DH5α carrying *mcr-1* strains against colistin and polymyxin B could appear similar changes. Venn diagram of DEPs in *E. coli* DH5α (pUC19-*mcr-1*) under different selection pressure of colistin and polymyxin B was shown in Fig 1C. Up to 16 DEPs including membrane protein (OmpX, RbsB, OmpT, and YtfJ), metabolic enzymes (Maa, IscU, YbiV, TreF, ThiG, AzoR, MsrC, and GcvH), DNA repair (DinG) and uncharactized protein (YdcH, YdeI, YbgS) were appeared simultaneously in CST group (CST0.5 and CST1) and PMB group (PMB0.5 and PMB1). GO analysis of DEPs whose abundance increases at different culture conditions showed significant enrichment of categories consistent with the key biological events related with antibiotic resistance. Observed categories such as metabolic process, biosynthesis of secondary metabolites and biosynthesis of amino acids (Fig 1D). Then we compared the variation trend of DEPs under the selection pressure of colistin and polymyxin B. Under the condition of colistin, among the total of 57 DEPs, 26 protein showed consistent changes in expression, of which 11 were down-regulated and 15 were up-regulated. Under the different concentration of polymyxin B, a total of 22 proteins in 57 DEPs showed consistent changes, of which 10 were down-regulated and 12 were up-regulated. Of the proteins with the same changes, 14 proteins appeared simultaneously under different selection pressures of colistin and polymyxin B, of which 5 proteins (TolA, YqaB, NadC, YbiG, Htp) were down-regulated and 9 proteins (GlpQ, OsmY, WaaJ, YaaF, EutE, YdcH, YjiY, GutD, and YccX) were up-regulated.

**Fig 1.**
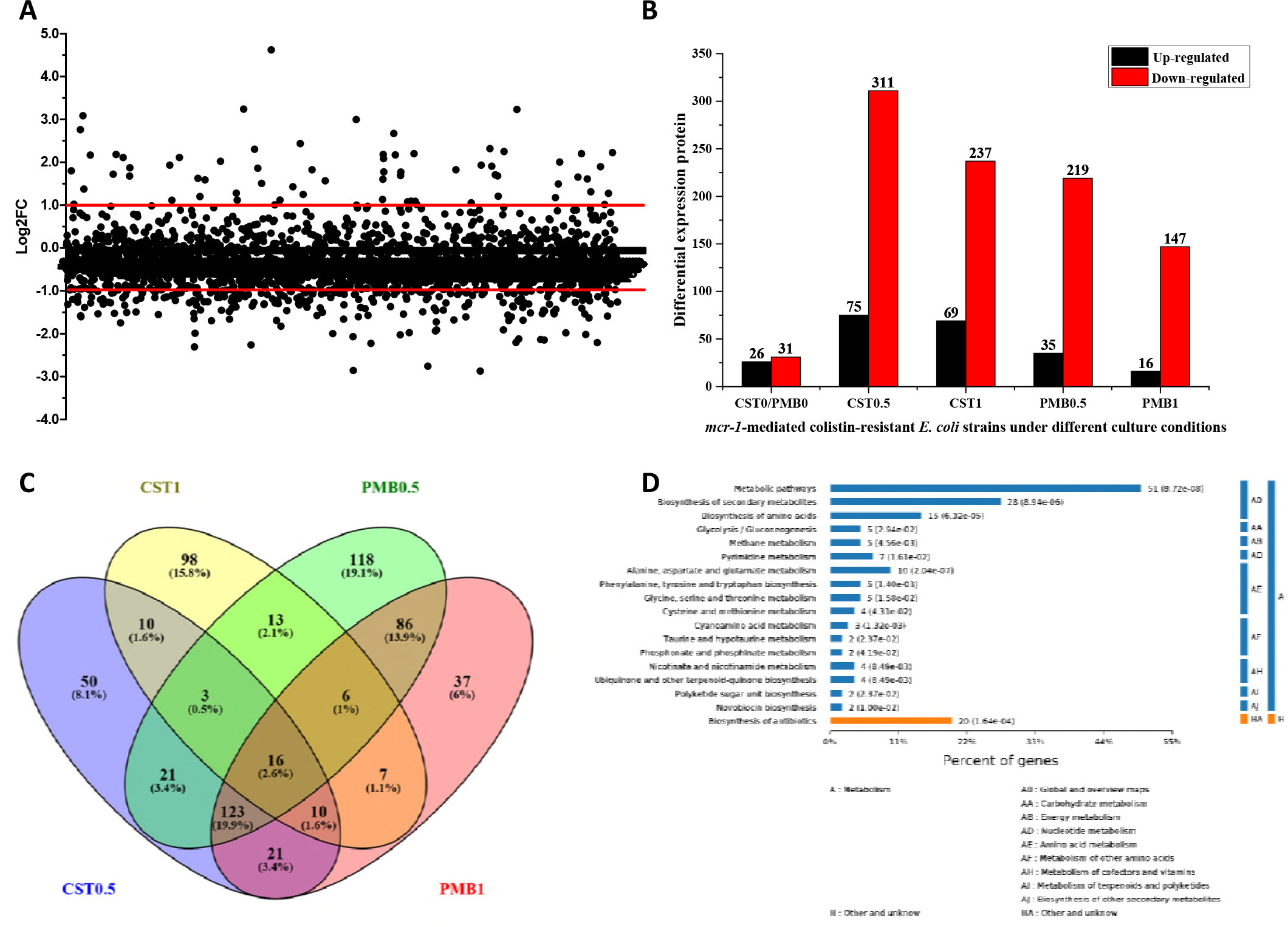
Comprehensive proteome profiles induced by *mcr-1* gene using label-free quantitative proteomics. (A) Protein distribution detected in colistin-resistant *E. coli* DH5α (pUC19-*mcr-1*) proteome using SWATH-MS. (B) Bar diagrams reflecting numbers of up-regulated and down-regulated proteins and in *E. coli* DH5α (pUC19-*mcr-1*) under different selection pressure of colistin and polymyxin B (no drug group (CST0/PMB0), colistin group (CST0.5 and CST1), and polymyxin B group (PMB0.5 and PMB1). (C) Venn diagram of DEPs in *E. coli* DH5α (pUC19-*mcr-1*) under different selection pressure of colistin and polymyxin B. (D) Metabolic pathways enriched with DEPs identified in colistin-resistant *E. coli* DH5α (pUC19-*mcr-1*) strains without the addition of drug.

Then we investigated the gene expression of these 14 consistent DEPs using qRT-PCR. The relative expression of the selected DEPs were shown in Fig 2. The gene expression of *htp*, *nadC*, *tolA*, *ybiG* and *yqaB* were down-regulated and were consistent with the proteomic analysis. The gene expression of *tolA* in colistin-resistant *E. coli* DH5α (pUC19-*mcr-1*) cultured in colistin group (CST0, CST0.5 and CST1) decreased by 0.08, 0.20, and 0.19-fold compared with that in colistin-sensitive *E. coli* DH5α (pUC19). Furthermore, the relative expression of *yjiY*, *eutE*, *glpQ*, *gutD*, *osmY*, *waaJ*, *yaaF*, *yccX*, and *ydcH* were significantly up-regulated in *E. coli* DH5α (pUC19-*mcr-1*). Noteworthy, *ydcH*, *osmY*, and *tolA* had the most significant changes in gene expression and it demonstrated that these three proteins might play an important role in *mcr-1*-mediated drug resistance.

**Fig 2.**
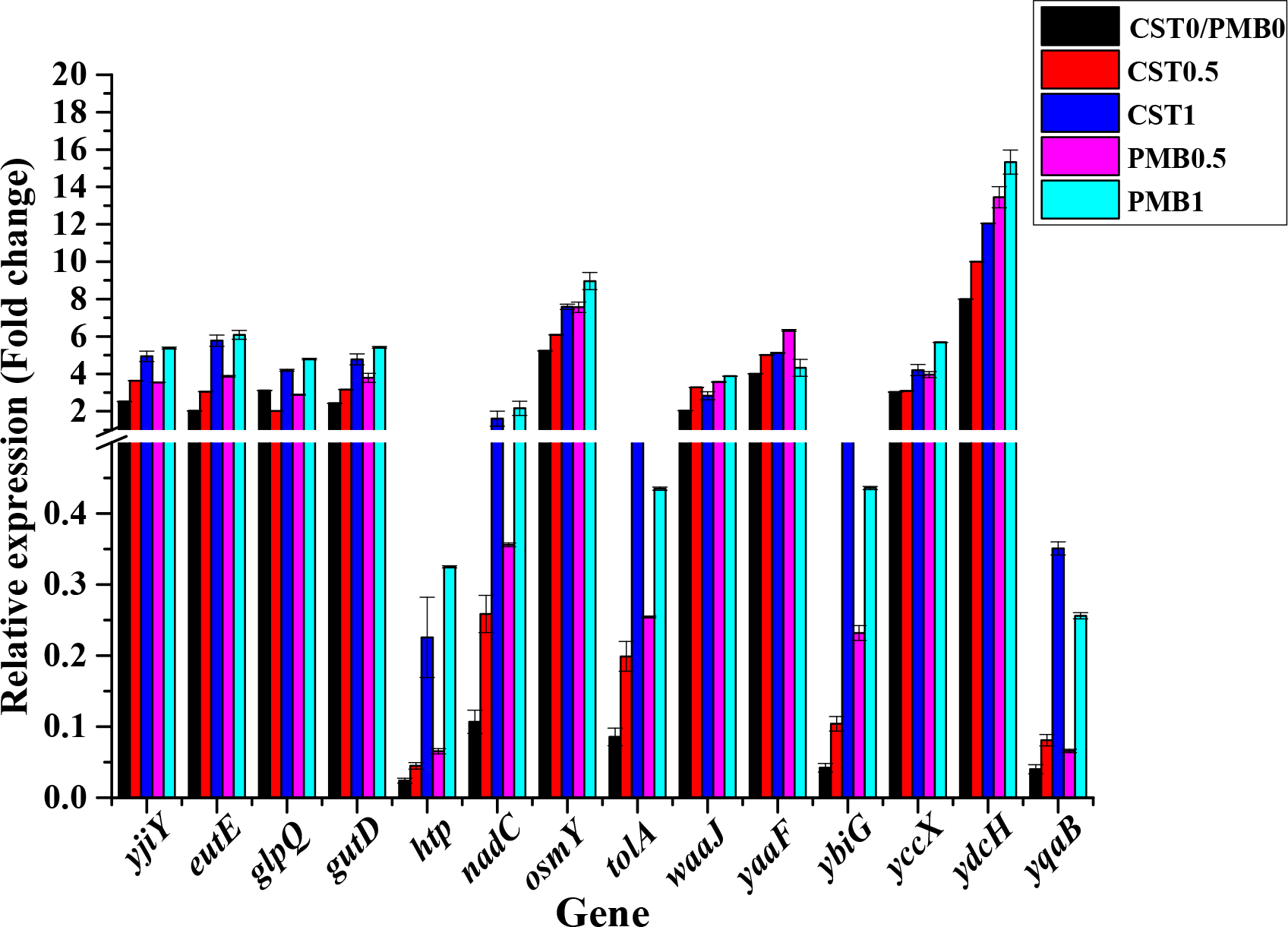
Transcript levels of a total of 14 proteins appeared simultaneously under different selection pressures of colistin and polymyxin. 16S rRNA was used as an internal control. All samples were prepared in three biological replicates. Error bars represented SEM. The data are normalized to the sensitive strain *E. coli* DH5α (pUC19).

### Functional analysis of proteins changed during antibiotic pressure

We expected that proteins changing in levels during drug selection pressure would play an important role in bacterial resistance. Pathway analysis of DEPs found that lysine biosynthesis, pyruvate metabolism, peptidoglycan biosynthesis were disturbed in colistin-resistant *E. coli* DH5α (pUC19-*mcr-1*) strains when no drug supplemented in the media. The further analysis found that the number of proteins involved in lipopolysaccharide biosynthesis was increased (up to five) when 0.5 *μ*g/mL of colistin was added. Whereas the amount of proteins involved in the synthesis of LPS decrease as the drug concentration continued to increase by a gradient. Similar changes could also be confirmed in the cationic polypeptide resistance known protein, peptidoglycan biosynthesis and glycerophospholipid metabolic pathway. Similarly, changes in thiamine metabolism, phenylalanine, tyrosine and tryptophan biosynthesis, and pyruvate metabolic processes were also largely affected (Fig 3). It was noteworthy that the expression of GlpQ, EutB, PgpC, PlsB, cdh, UgpQ, aas, EutC, and GlpC involved in glycerophospholipid metabolism were up-regulated in colistin-resistant *E. coli* DH5α (pUC19-*mcr-1*) under the selection pressure of colistin.

**Fig 3.**
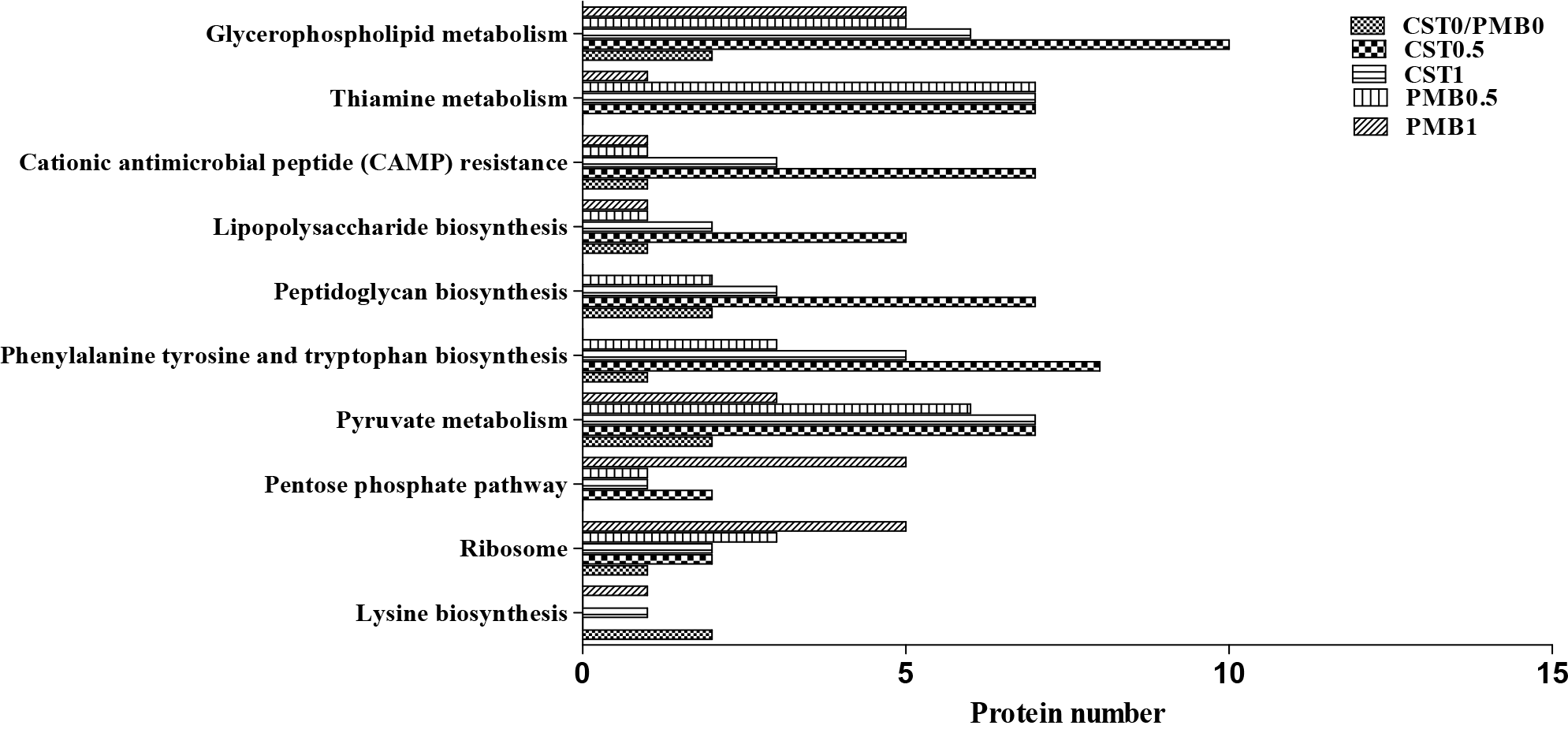
KEGG pathway analysis of DEPs in *E. coli* DH5α (pUC19-*mcr-1*) strains under different selection pressures of colistin and polymyxin B.

Under different concentrations of polymyxin B selection pressure, DEPs showed the same trend of expression, and the proteins involved in the metabolic processes varied greatly. These metabolic processes mainly involved in glycerophospholipid metabolism, thiamine metabolism, lipopolysaccharide biosynthesis, and cationic antibacterial peptide (CAMP) resistance (Fig 3). However, there was no significant change in the relevant proteins involved in CAMP resistance after the addition of polymyxin B to the culture medium, unlike colistin. In addition, *E. coli* DH5α (pUC19-*mcr-1*) strains in different concentrations of polymyxin B selection pressure, with the drug concentration increasing, the number of affecting the function of ribosomal protein was increased, which indicating that the protein synthesis process has been affected. The bacteria were bound to enhance protein synthesis in order to adapt to drug selection pressure. In terms of energy metabolism, part of the protein involved in the TCA cycle and pentose phosphate pathway was up-regulated, indicating an increase in energy metabolism to accommodate drug stress (Fig 3).

### Untargeted metabolomic analysis

Untargeted metabolomic profiling of colistin-resistant *E. coli* DH5α (pUC19-*mcr-1*) and colistin-sensitive *E. coli* DH5α (pUC19) strains under different culture conditions was studied to obtain their metabolic characteristics. First, PCA was used to investigate the metabolomic profile of the two strains cultured in different concentrations of colistin. The PCA analysis separated on BEH RP C_18_ chromatographic column in ESI^+^ and ESI^−^ mode were shown in Fig 4A and 4C, respectively. The intracellular metabolites between the colistin-resistant and -sensitive strains cultured in different drug concentrations were significantly different. Clustering analysis of any pair of comparisons in their respective space was obvious. Then orthogonal projection to latent structures-discriminant analysis (OPLS-DA) were applied to identify the main difference variables. S-plot of colistin-resistant and - sensitive strains separated on BEH RP C_18_ column in ESI^+^ and ESI^−^ mode were shown in Fig 4B and 4D, respectively. A total of 67 differential metabolites were simultaneously found in different drug culture conditions, of which 49 metabolites were selected in ESI^+^ mode, 18 metabolites were found in ESI^−^mode (Table 1). Among all the identified metabolites separated on C_18_ column, 24 features (10 up-regulated and 1 down-regulated) were found in ESI^+^ mode and 13 features (10 up-regulated and 3 down-regulated) were selected in ESI^−^mode, respectively. Then we analyzed the metabolites separated on HILIC column. A total of 38 compounds (18 up-regulated and 20 down-regulated) were identified in ESI^+^ mode and 5 features (4 up-regulated and 1 down-regulated) were selected in ESI^−^ mode, respectively (Table 1). For QC purposes, several reference standards were added to verify the reliability of the metabolomics analysis. The average error of calculated mass was below 10 ppm, the retention time was nearly identical, and the RSD of the peak area in three batches (different extracts) was 3.52%-6.24% (Table S7).

**Table 1.**
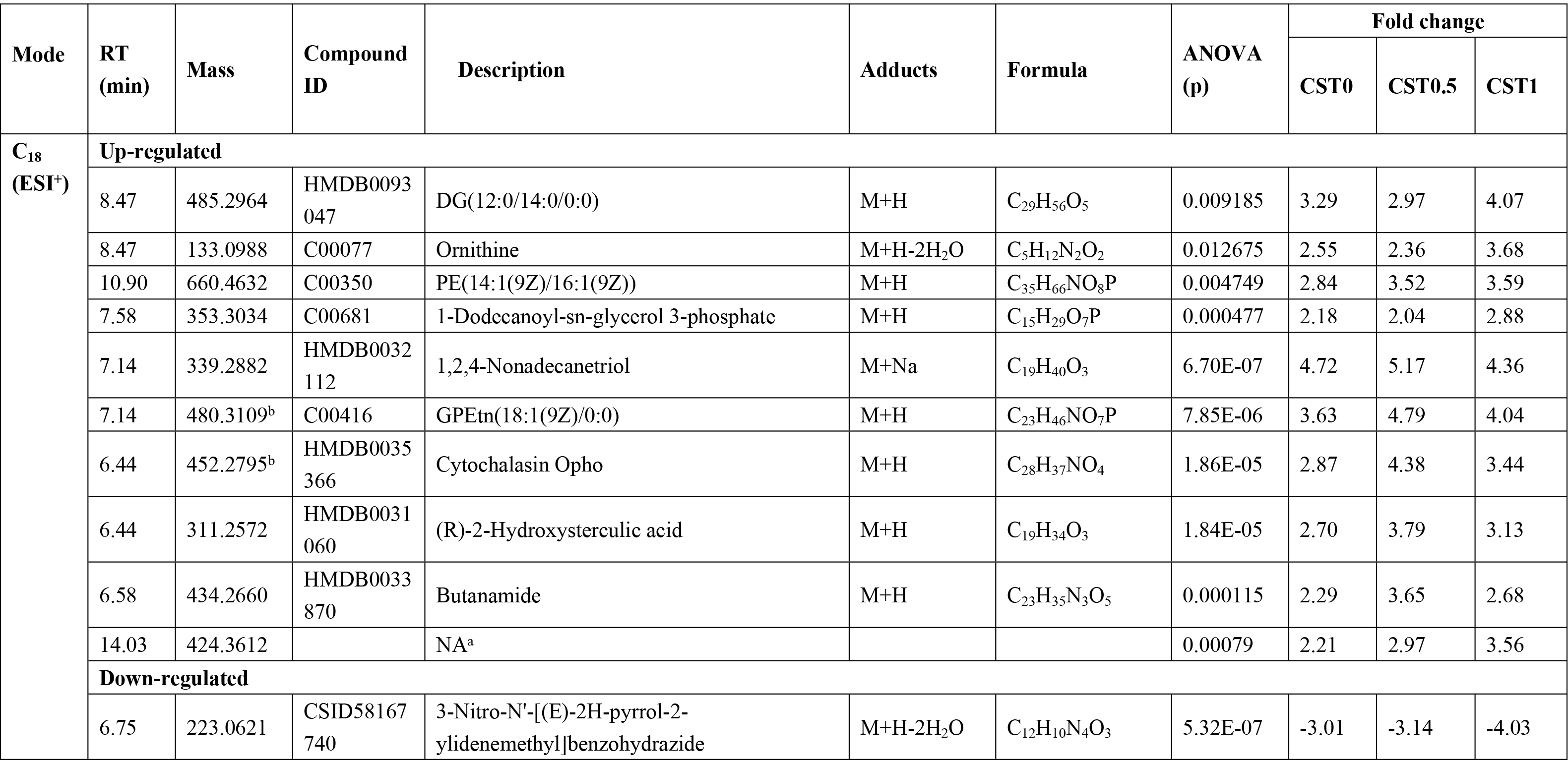
Putative differential metabolites identified in colistin-resistant *E. coli* DH5α (pUC19-*mcr-1*) under three different drug culture conditions.

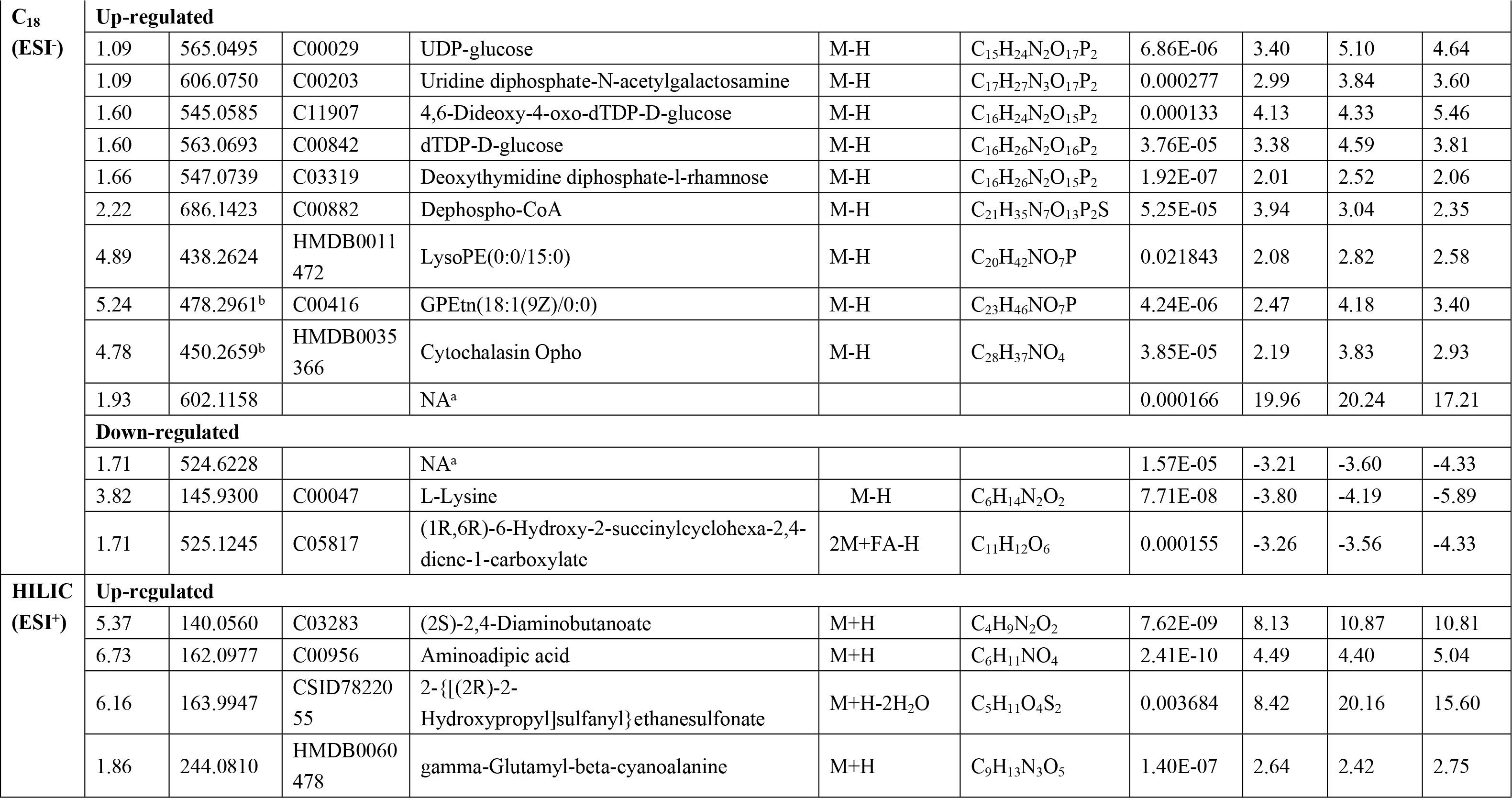

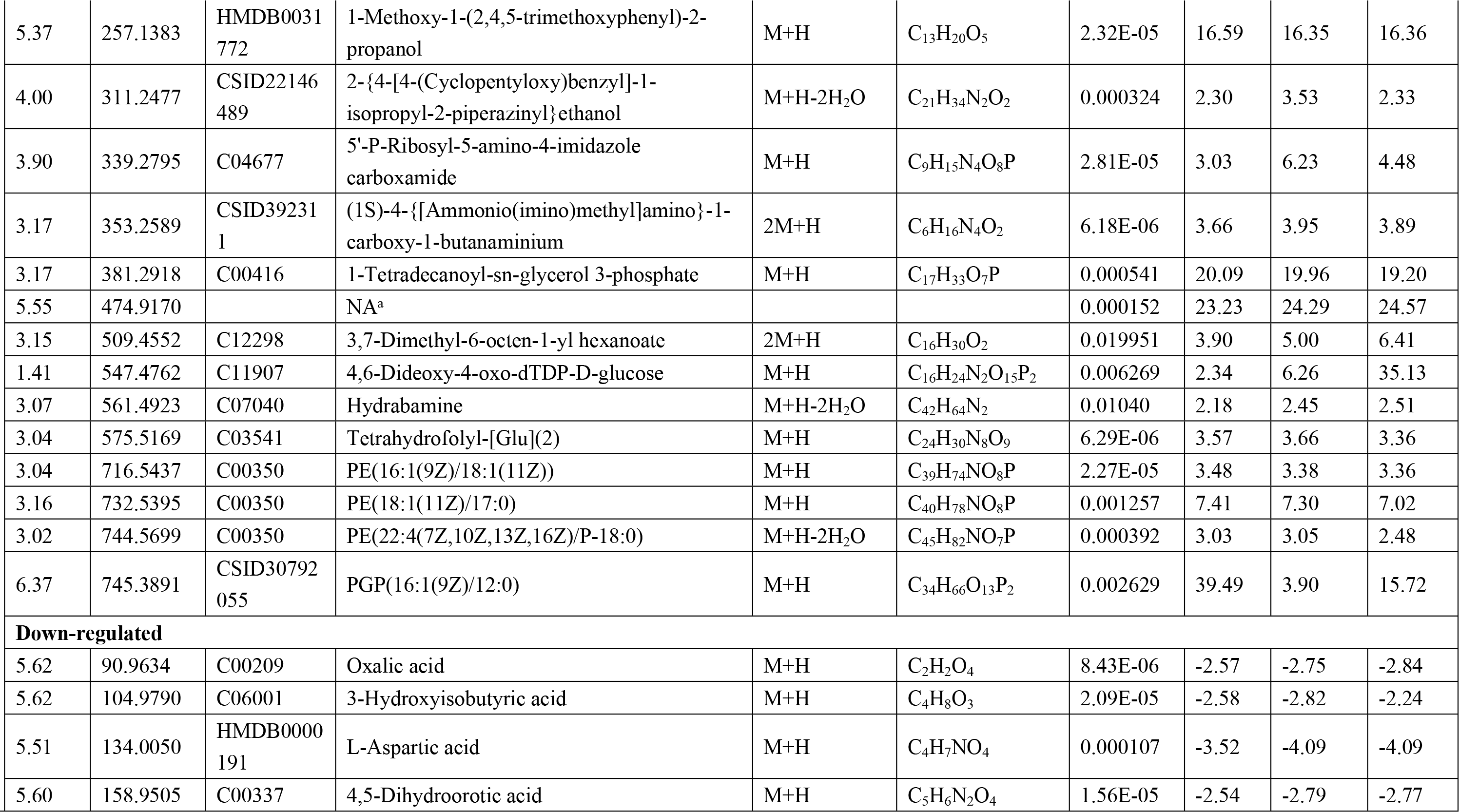

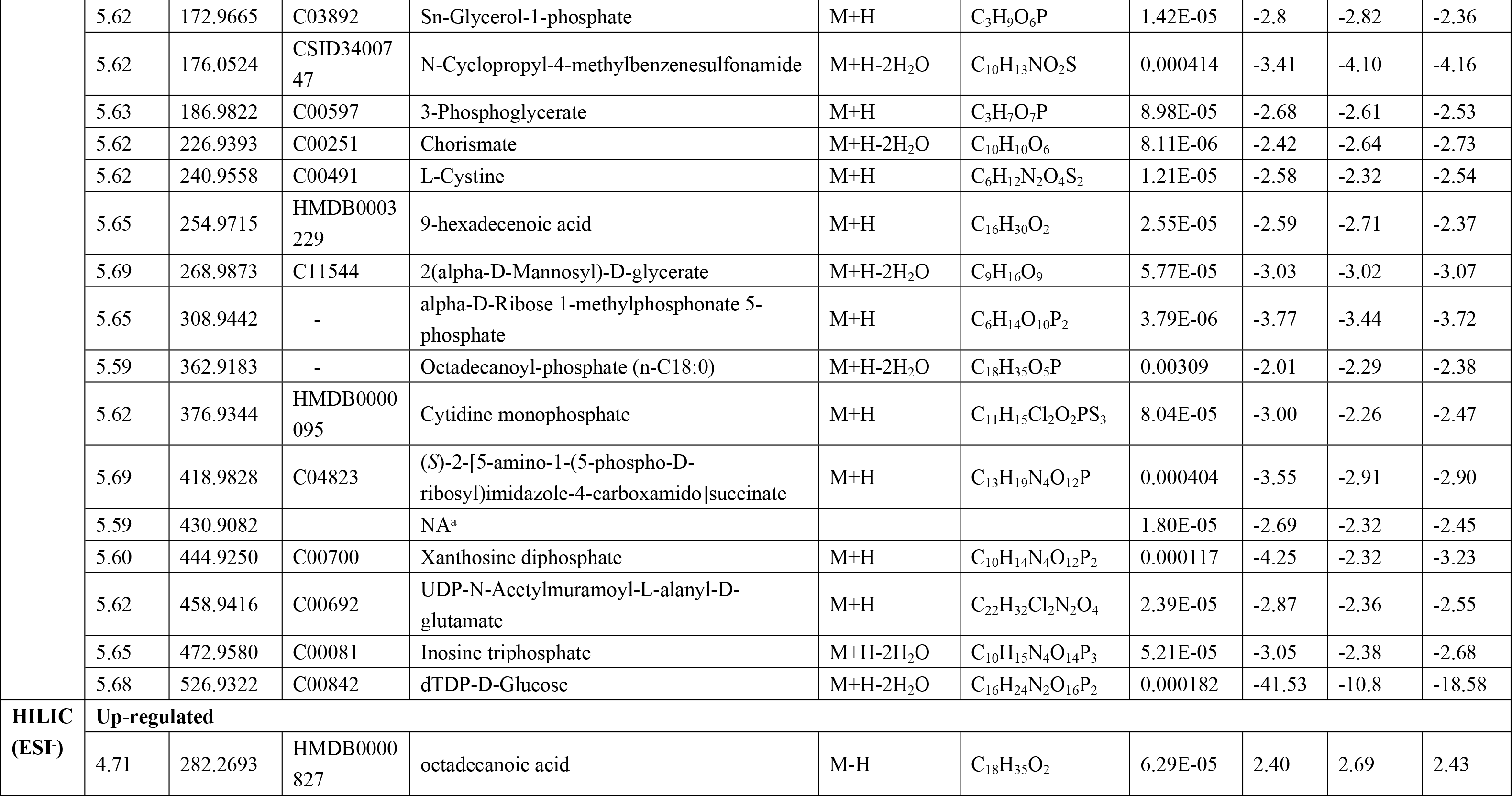

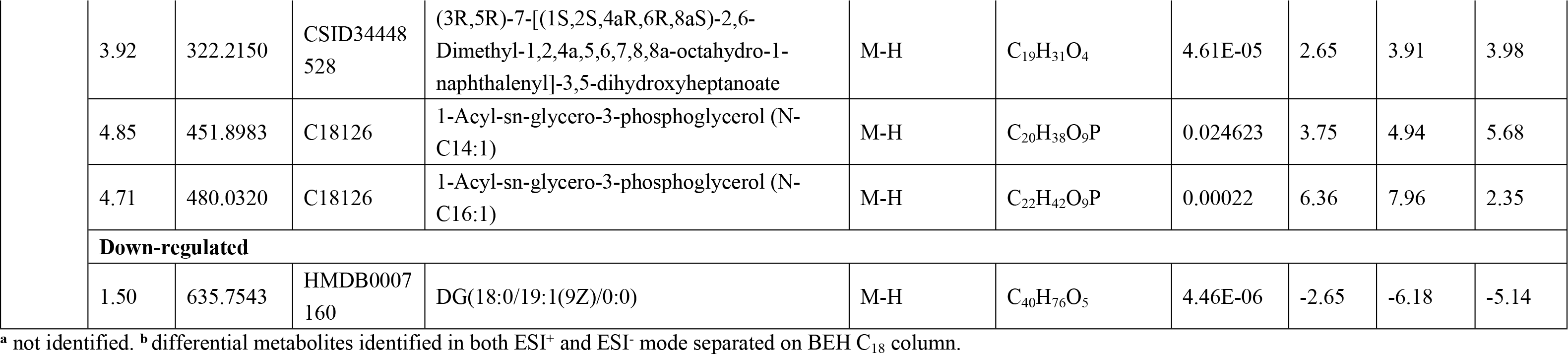

**Fig 4.**
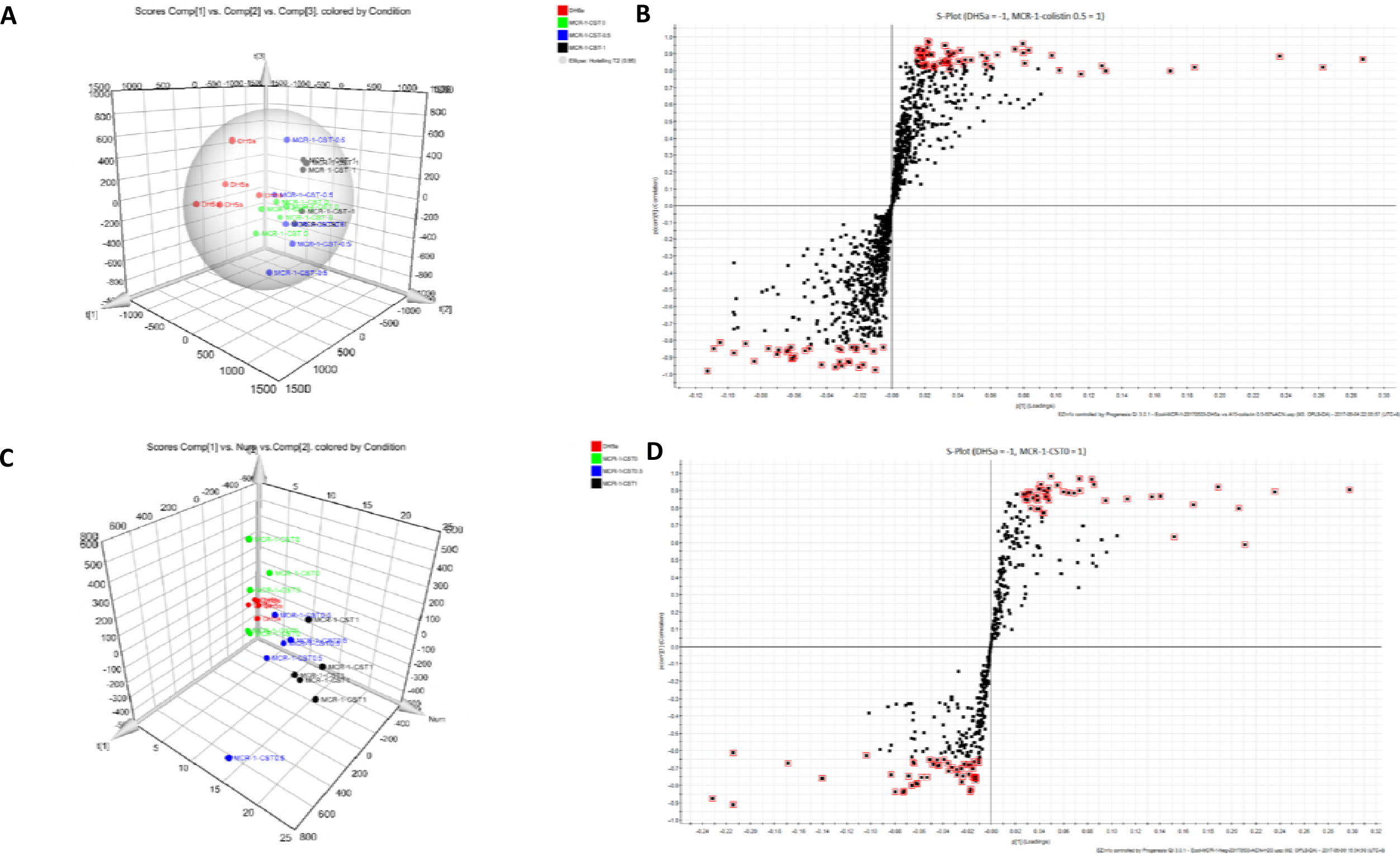
Metabolic characteristic of the cellular metabolites of *E. coli* sensitive and resistant strains under different concentrations of colistin separated on BEH C_18_ column. (A) Principal component analysis of microbial extracts from resistant and sensitive stains in ESI+ mode. (B) Representative S-plot of resistant strains versus sensitive strains in ESI+ mode. (C) Principal component analysis of microbial extracts from resistant and sensitive stains in ESI^−^ mode. (D) Representative S-plot of resistant strains versus sensitive strains in ESI^-^ mode.

### Enrichment and metabolic pathway analysis

Heatmap presentation of all identified intracellular differential metabolites from colistin culture group (CST0, CST0.5 and CST1) was shown in Fig 5A. To facilitate access to metabolic pathway analysis, altered metabolites from the merged data set were mapped to KEGG and obtained the disturbed metabolic pathway using MetaboAnalyst 3.0. Metabolite set enrichment analysis showed that pantothenate and CoA biosynthesis, glycerolipid metabolism, phosphatidylethanolamine biosynthesis were contributed to *mcr-1* mediated bacterial metabolism (Fig 5B). Then we carried out the metabolic pathway analysis and found that polyketide sugar unit biosynthesis, glycerophospholipid metabolism, and lysine biosynthesis were significantly enriched for all the identified differential metabolites (Fig 5C). Interestingly, protein-metabolite interaction analysis of the disturbed glycerophospholipid metabolism in *mcr-1*-mediated colistin resistance by the integration of proteomics and metabolomics found that a high number of differential metabolites including PE (14:1(9Z)/16:1(9Z)), GPEtn (18:1(9Z)/0:0), LysoPE (0:0/15:0), GPEtn (18:1(9Z)/0:0), PE (16:1(9Z)/18:1(11Z)), PE (18:1(11Z)/17:0), PE (22:4(7Z,10Z,13Z,16Z)/P-18:0), PGP (16:1(9Z)/12:0), sn-glycerol-1-phosphate, 1-acyl-sn-glycero-3-phosphoglycerol (N-C14:1), 1-acyl-sn-glycero-3-phosphoglycerol (N-C16:1) involved in glycerophospholipid metabolism were up-regulated in the colistin-resistant *E. coli* DH5α (pUC19-*mcr-1*) strains compared with that in the colistin-sensitive *E. coli* DH5α (pUC19) strains (Fig 5D).

**Fig 5.**
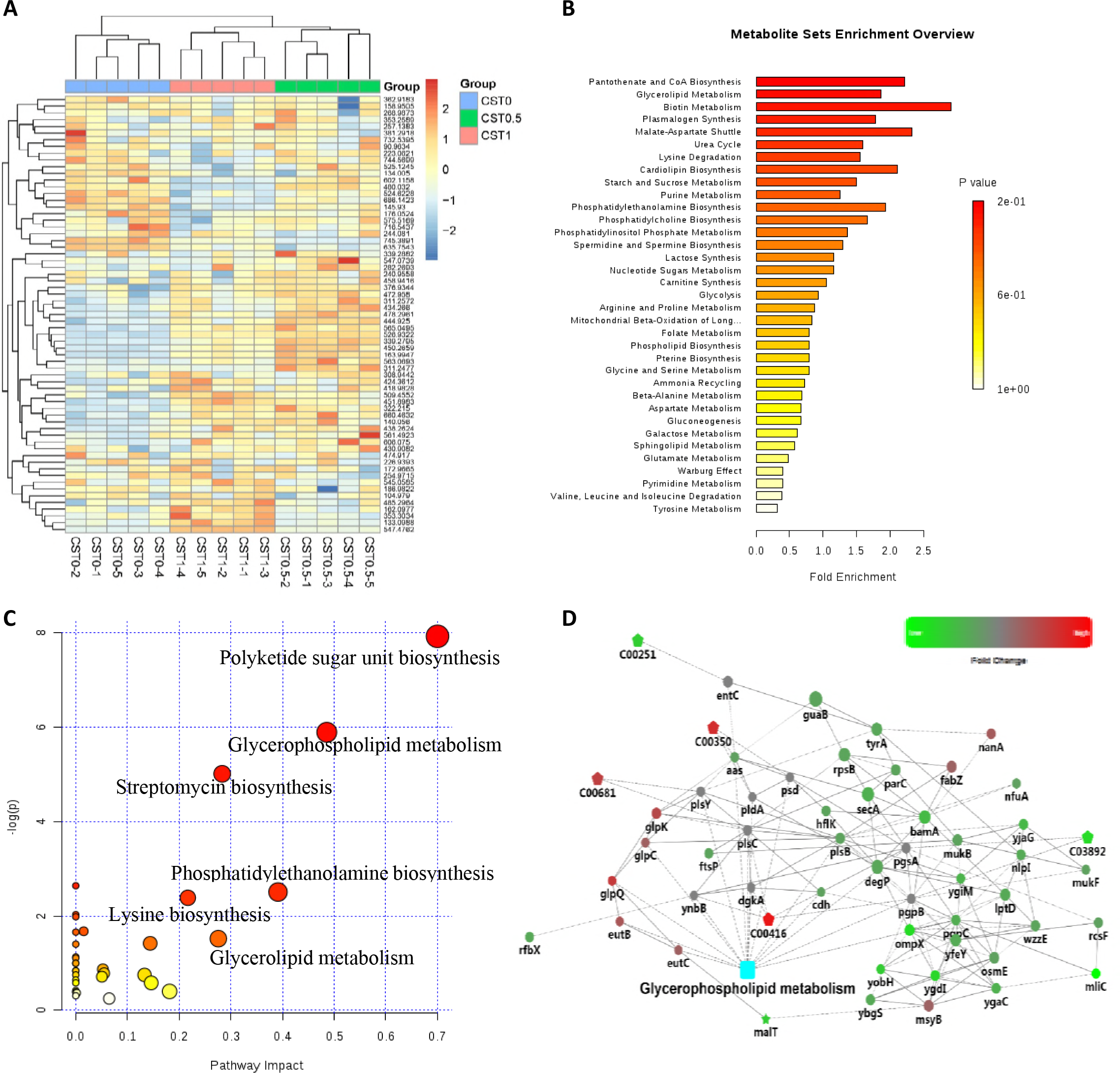
Enrichment and metabolic pathway analysis of *mcr-1*-mediated colistin resistance using untargeted metabolomics. (A) Heatmap presentation of intracellular metabolic profiles from *E. coli* DH5α (pUC19-*mcr-1*) strains under different concentrations of colistin culture conditions (CST0, CST0.5 and CST1). Each column represents one biological replicate, and each row represents one targeted metabolite detected in this study. (B) Metabolite set enrichment analysis showed that pantothenate and CoA biosynthesis, glycerophospholipid metabolism, glycerolipid metabolism, phosphatidylethanolamine biosynthesis were contributed to *mcr-1* mediated bacterial disturbed metabolism. (C) Visualization of disturbed metabolic pathways in *mcr-1* mediated colistin-resistant strains. (D) Protein-metabolite interaction analysis of the disturbed glycerophospholipid metabolism in *mcr-1*-mediated colistin resistance by the integration of proteomics and metabolomics. Rectangular nodes represent KEGG pathway; Round nodes indicate protein; Pentagon represents metabolites. Red: up-regulated; Green: down-regulated.

## Discussion

At present, it was preliminarily thought that the resistant mechanism of *mcr-1* in colistin resistance was that the inner membrane protein MCR-1 was chemically modified as a form of PEA on the lipid A group of bacterial LPS [27–29]. As a lipid-modified gene, whether *mcr-1* brings comprehensive proteomic and metabolomic changes in the bacteria and affects the corresponding metabolic pathway is largely unknown. This study is the first to investigate the comprehensive proteomic and metabolomic profiles of *mcr-1*-mediated colistin-resistant and -sensitive *E. coli* strains under different selection pressures of colistin and polymyxin B.

It is well established that polymyxins exert their antimicrobial activity mainly through the disruption of the bacterial outer membrane [30, 31]. We used a label-free quantitative proteomics approach to elucidate the protein profiles of colistin-resistant *E. coli* DH5α carrying *mcr-1* under different drug selection pressure, and compared them with the expressed proteome in the colistin-sensitive sample. Our findings demonstrated colistin and polymyxin B made *mcr-1*-mediated resistant bacteria to exhibit different proteome profiles and it suggest that colistin-resistant bacteria might alter their metabolism to adapt to antibiotic resistance (Fig 1D). Proteomic characteristics discovered by SWATH-MS showed that *mcr-1* could cause the disturbed metabolic regulation such as glycerophospholipid metabolism, thiamine metabolism, LPS biosynthesis, and CAMP resistance (Fig 3) and the protein profile of *E. coli* carrying *mcr-1* to colistin showed the most significant changes, which was consistent with the report of Hua et al [24]. Integrative proteomic and metabolomic analysis demonstrated that GlpQ, EutE involved in glycerphospholipid metabolism were significantly up-regulated in colistin-resistant *E. coli* DH5α (pUC19-*mcr-1*) under the condition of blank LB broth. However, the number of DEPs (GlpQ, EutB, PgpC, PlsB, cdh, UgpQ, aas, EutC, and GlpC) involved in glycerphospholipid metabolism increased when the resistant *E. coli* DH5α (pUC19-*mcr-1*) cultured in the different drug conditions. In addition, metabolite expression analysis found that the metabolites involved in glycerophospholipid metabolism were up-regulated due to the insertion of *mcr-1*. Remarkably, the substrate phosphatidylethanolamine (Kegg ID: C00350) for *mcr-1* to mediate colistin resistance was investigated as the differential metabolites and its expression in different colistin selection pressure (CST0, CST0.5, and CST1) were up-regulated in resistant *E. coli* DH5α (pUC19-*mcr-1*) strains. It was predicted that the accumulation of phosphatidylethanolamine was appeared in *mcr-1*-mediated colistin resistance in *E. coli*. Glycerophospholipids are important membrane lipids in bacteria and play a vital role in cellular functions, including the regulation of transport processes, protein function, and signal transduction [32]. Additionally, glycerophospholipids are essential components of lipoproteins and influence their function and metabolism [33]. These results demonstrated that *mcr-1* could cause disturbed glycerophospholipid metabolism pathway to adapt colistin selection pressure in *E. coli*.

In addition of the effect on the membrane lipids, *mcr-1* caused metabolic perturbation of LPS biosynthesis in colistin-resistant strains. LPS is a key component of the outer membrane, a permeability barrier in Gram-negative bacteria, in which it plays important roles in the integrity of the outer-membrane permeability barrier [34, 35]. When the *E. coli* strains carrying *mcr-1* were cultured under the selection pressure of colistin and polymyxin B, seven members (LpxC, KdsA, KdtA, RfaI, RfaJ, lpp and LptB) of LPS transport protein family together to transport LPS from the inner membrane to the outer leaflet of the outer membrane and the expression of these seven members were decreased in *mcr-1*-mediated colistin-resistant *E. coli*. So we predicted that *mcr-1* might influence the biosynthesis and transport of lipoprotein in colistin resistance.

Furthermore, colistin and polymyxin B also brought changes in the relevant efflux proteins involved in CAMP resistance (kegg ID: ko01503). Efflux pumps could form multicomponent “pumps” that span both inner and outer membranes in Gram-negative bacteria and confer resistance to a broad spectrum of antibiotics [36, 37]. The expression of efflux pump components was jointly controlled by several positive global regulators, which respectively decrease and enhance the transcription of specific genes, such as acrAB-tolC [38]. Our findings indicated that a higher number of membrane protein TolC, NlpE, AcrA, AcrB, SapD, depG, and TolA had changed under the concentration of colistin as 0.5 *μ*g/mL, compared with the number of DEPs in the condition of colistin of 1 *μ*g/mL. The insertion of *mcr-1* in *E. coli* not only caused the PEA modification of bacterial cell membrane lipid A, but also affected the efflux of the polymyxins through disturbing the expression of efflux pump proteins involved in CAMP resistance pathway.

In addition, the number of the DEPs that affecting the ribosomal function were increased in *E. coli* DH5α (pUC19-*mcr-1*) strains with the increasing of exposed polymyxin B concentration and it indicated that the protein synthesis process has been disturbed. The bacteria could enhance protein synthesis in order to adapt to drug
selection pressure. In terms of energy metabolism, parts of the protein involved in the TCA cycle and pentose phosphate pathway was upregulated, indicating an increase in energy metabolism to accommodate drug stress. Increased metabolic activity and energy production are required for pathogens to resist high antibiotics stress [39]. The metabolic changes may be a consequence of the biological cost of antibiotic resistance. Lysine biosynthesis was critical for protein biosynthesis and a component of the peptidoglycan layer of bacterial cell walls [40]. Metabolomic analysis found that the expression of lysine and L-2-aminoadipate was down-regulated in colistin-resistant *E. coli* DH5α (pUC19-*mcr-1*) compared with the sensitive strains.

In conclusion, our data highlight the comprehensive proteome and metabolome profiles of *mcr-1*-mediated colistin resistance in *E. coli* and different proteome characteristics are exhibited under different selection pressure of colistin and polymyxin B. The substrate phosphatidylethanolamine for *mcr-1* to mediate colistin resistance is accumulated in colistin-resistant *E. coli*. Notably, *mcr-1* can not only cause the phosphoethanolamine modification of bacterial cell membrane lipid A, but also affect the efflux of the colistin through disturbing the expression of efflux pump proteins involved in cationic antibacterial peptide resistance pathway. Furthermore, the disturbed glycerophospholipid metabolism is closely related with *mcr-1*-mediated colistin resistance and this finding can further provide valuable information to inhibit colistin resistance by blocking this metabolic process.

## Supporting Information

**Table S1** Primers used in qRT-PCR analysis.

**Table S2** Identification of the DEPs in colistin-resistant *E. coli* DH5α (pUC19-*mcr-1*) strains cultured in blank media.

**Table S3** DEPs selected in resistant *E. coli* DH5α (pUC19-*mcr-1*) cultured in the concentration of polymyxin B of 0.5 *μ*g/mL.

**Table S4** DEPs selected in resistant *E. coli* DH5α (pUC19-*mcr-1*) cultured in the concentration of polymyxin B of 1 *μ*g/mL.

**Table S5** DEPs selected in resistant *E. coli* DH5α (pUC19-*mcr-1*) cultured in the concentration of colistin of 0.5 *μ*g/mL.

**Table S6** DEPs selected in resistant *E. coli* DH5α (pUC19-*mcr-1*) cultured in the concentration of colistin of 1 *μ*g/mL.

**Table S7** Mass quality control results of UPLC-QTOF/MS untargeted metabolomics analysis.

